# Subcallosal area 25: its responsivity to the stress hormone cortisol and its opposing effects on appetitive motivation in marmosets

**DOI:** 10.1101/2023.12.11.571112

**Authors:** Rana Banai Tizkar, Lauren McIver, Christian Michael Wood, Angela Charlotte Roberts

## Abstract

Aberrant activity in caudal subcallosal anterior cingulate cortex (scACC) is implicated in depression and anxiety symptomatology, with its normalisation a putative biomarker of successful treatment response. The function of scACC in emotion processing and mental health is not fully understood despite its known influence on stress-mediated processes through its rich expression of mineralocorticoid and glucocorticoid receptors. Here we examine the causal interaction between area 25 within scACC (scACC-25) and the stress hormone, cortisol, in the context of anhedonia and anxiety-like behaviour. Moreover, the overall role of scACC-25 in hedonic capacity and motivation is investigated under transient pharmacological inactivation and overactivation. The results suggest that a local increase of cortisol in scACC-25 shows a rapid induction of anticipatory anhedonia and increased responsiveness to uncertain threat. Separate inactivation and overactivation of scACC-25 increased and decreased motivation and hedonic capacity, respectively, likely through different underlying mechanisms. Together, these data show that area scACC-25 has a causal role in consummatory and motivational behaviour and produces rapid responses to the stress hormone cortisol, that mediates anhedonia and anxiety-like behaviour.

## Introduction

One brain region repeatedly reported to be overactive in depression and anxiety is the caudal subcallosal cingulate cortex[1,2] (scACC). Consistent with this is the finding that reduction of activity within this area, regardless of treatment modality[3–5], coincides with successful outcomes. Moreover, intervention studies in marmoset monkeys provide evidence for the causal role of overactivation of area 25 within caudal scACC (scACC-25) in the symptoms of depression and anxiety. Specifically, scACC-25 overactivation induces anticipatory and motivational anhedonia[6], a major symptom of depression and especially common in treatment resistant patients[7,8]. It also heightens reactivity to uncertain threat, causes generalisation of conditioned threat responses and increases sympathetic cardiovascular activity alongside reduced heart rate variability[9]; symptoms commonly reported in generalised anxiety disorder[10,11]. This raises the question, what natural conditions or contexts is scACC-25 activated to heighten reactivity to threat and blunt anticipatory and motivational approach responses?

One likely candidate is stress. For example, area 25/14 scACC activity has been shown to correlate positively with plasma levels of the stress hormone, cortisol, regardless of behavioural context in macaque monkeys[12]. Moreover, scACC-25 expresses high levels of both mineralocorticoid (MR) and glucocorticoid (GR) receptors[13] making it sensitive to stress-mediated effects including circulating cortisol. However, the relationship of scACC-25 with stress is likely to be a complex one, since not only is scACC-25 responsive to cortisol but through its projections onto the hypothalamus[14] can regulate levels of cortisol release. To address this issue, we determined the effects of infusing cortisol directly into scACC-25. It was predicted that cortisol may induce a similar effect in scACC-25 as overactivation, consistent with the hypothesis that stress induces scACC-25 activation. Accordingly, we assessed its effects on both reactivity to uncertain threat, as measured with the human intruder paradigm and anticipatory appetitive arousal as measured by Pavlovian, discriminative conditioning (experiment 1).

The second issue this study was designed to address was the relative contributions of scACC-25 to the regulation of appetitive and threat-related behaviours. Previously, Wallis and colleagues[15] showed that inactivation of scACC-25 in marmosets reduced conditioned threat responses, enhanced parasympathetic cardiovascular activity and increased heart rate variability, opposing effects to that seen following overactivation[9]. In contrast, inactivation appeared to have no impact on responsivity to rewarding stimuli as measured by Pavlovian conditioned appetitive responses[6], suggesting limited basal activity during reward anticipation, despite the recent macaque data that indicates punctate scACC-25 firing during reward anticipation[16]. To determine whether scACC-25 is involved in the regulation of other aspects of appetitive responding beyond anticipatory arousal, we studied the effects of inactivation on appetitive motivation, as measured by a marmoset’s willingness to make progressively more responses for reward, preference and appetitive consumption, as measured by the sucrose preference test (experiment 2). For both studies marmosets received surgery to implant intracerebral cannulae targeting scACC-25, and following suitable recovery received localised scACC-25 microinfusions of varying concentrations of cortisol (experiment 1) and a cocktail of GABA_A_and GABA_B_receptor agonists for inactivation purposes and dihydrokainic acid (DHK, glutamate reuptake inhibitor) for overactivation purposes (experiment 2).

## Methods

### Subjects

Twelve marmosets (Callithrix jacchus, 6 females), bred on-site at the University of Cambridge Marmoset Breeding Colony, were housed in male/female pairs (males were vasectomized). Four subjects (2 females) were in Experiment 1 while eight (4 females) were in Experiment 2 (for details of each subject’s experience refer to Table S1). All procedures were carried out in accordance with the UK Animals (Scientific Procedures) Act 1986 and the University of Cambridge Animal Welfare and Ethical Review Body.

### Surgical procedures

All subjects received chronic cannulae implanted into scACC-25 at coordinates +14.0 AP; ±0.7 LM with in situ adjustments if required[17,18]. Four subjects (experiment 1) received telemetry probe implants for the transmission and recording of real-time cardiovascular activity. Marmosets were premedicated with ketamine hydrochloride (0.1mL of 100mg/mL solution, IM, Vetalar,) and the nonsteroidal anti-inflammatory analgesic meloxicam (0.075mL of 20mg/mL solution s.c., Metacam, Boehringer Ingelheim). For full details of surgical procedure, care and medication refer to Supplemental Methods.

### Drug treatments

Drugs were delivered bilaterally via sterile injectors (Plastic One, C235I/SPC). A pump delivered a constant rate of 0.5μl/min for DHK (6.25nmol/μl; Tocris, UK – pretreatment time of 10min) and two doses of cortisol (1.5 and 5ng/μl diluted in 0.9% saline; Hydrocortisone hemisuccinate, Sigma-Aldrich – pretreatment of 8min), and a rate of 0.25μl/min for the muscimol/baclofen cocktail (0.1mM muscimol/1.0mM baclofen; Sigma-Aldrich – pretreatment of 25min) for a total duration of 2 minutes. Subcutaneous injection of 20mg/kg cortisol (or 0.9% saline as control) was administered with a pre-treatment time of 15 minutes (summarised in Table S2).

### Behavioural testing apparatus

Both the progressive ratio and appetitive Pavlovian tests were carried out in a sound-attenuated apparatus, whilst the human intruder and sucrose preference tests were carried out in the top right quadrant of the home cage, where the subject was divided from the partner.

#### Human intruder test

The human intruder test was used to measure the intolerance of marmosets to uncertainty under experimental manipulations (described in detail in Quah and colleagues[19]). An experimenter wearing a realistic latex human mask (Greyland Film, UK) and standard lab attire entered the room and stood in front of the home cage for two minutes, maintaining constant eye contact with the subject (Intruder phase). Behaviour was scored offline (JWatcher software), measuring the time subjects spent in each zone (Figure 1A). Their location, percentage time moving, head and body bobs and vocalisations (recorded by a shotgun microphone) were compiled into an exploratory factor analysis (EFA) score (Additional details see Supplemental Methods and Figure S1). Increase in this score indicates an increase in anxiety-like behaviour.

**Figure 1.**
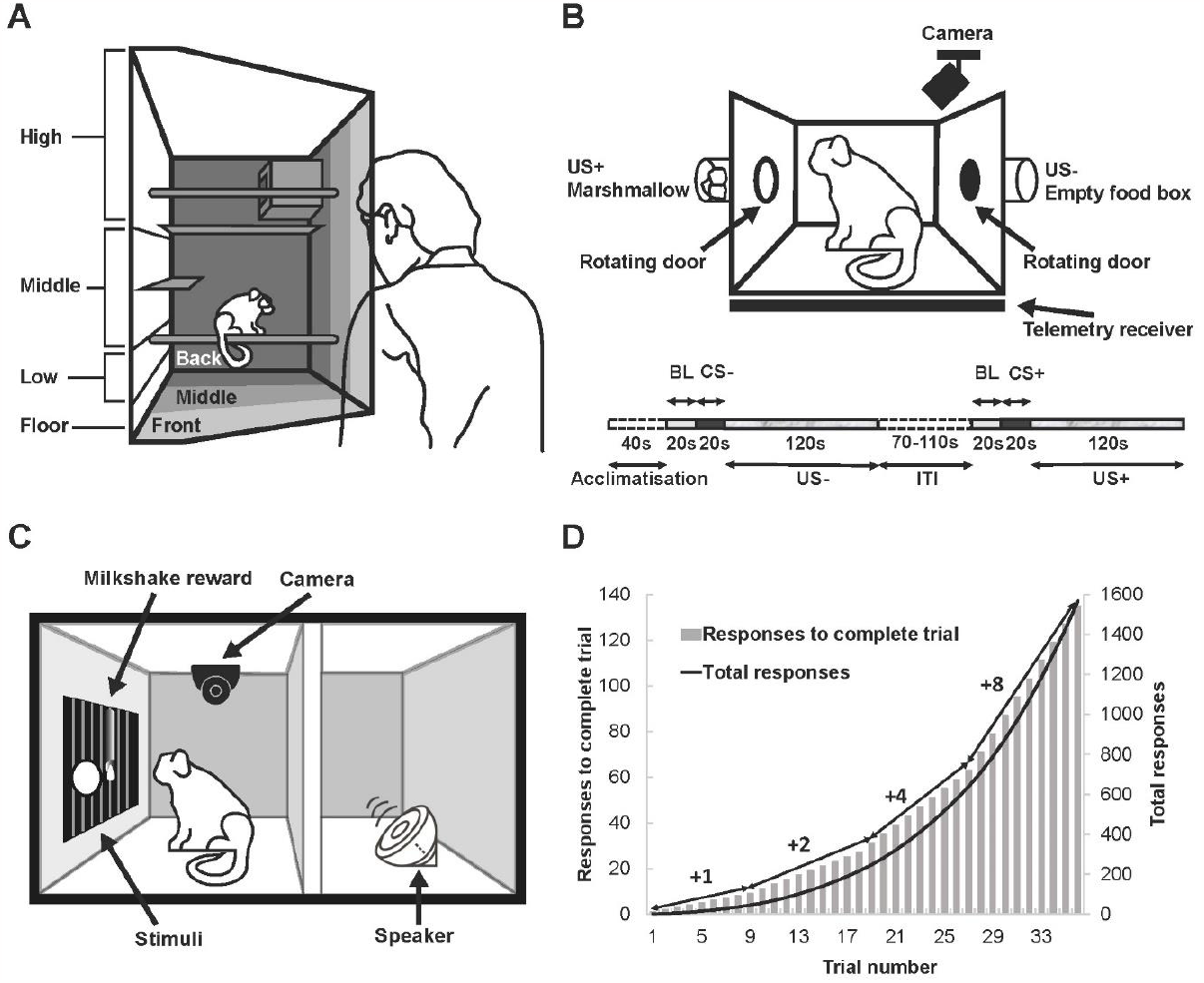
Behavioural paradigms. A. Human intruder test carried out in home cage illustrating zone categorisation. B. Schematic illustration of appetitive Pavlovian test apparatus (above) with a timeline of a two-trial session (below) used for all infusions. C. Schematic illustration of touchscreen-based test apparatus for progressive ratio test. D. Graph illustrating the progressive ratio schedule.

#### Appetitive Pavlovian test

This test consisted of two trial types, rewarded and non-rewarded. In rewarded trials, access to a food box containing marshmallows (Unconditioned Stimulus, US+) was provided for 120s after a neutral sound cuing the reward (Conditioned Stimulus, CS+) was played for 20s. In non-rewarded trials a neutral sound cued no reward (CS-) which was followed by display of an empty food box (US-) for 120s on the opposite side of the apparatus (details of training in supplemental methods). Experimental manipulations occurred shortly prior to sessions containing a non-rewarded trial followed by a rewarded trial (Figure 1B).

Anticipatory arousal for reward is measured by rapid head rotations from side to side, called head jerks[20], as well as raised mean arterial pressure (MAP). For analyses, CS-directed data were calculated as the relative change in these measures compared to the 20-second baseline (BL) period prior to the CS period. The US-directed responses were analysed relative to the CS period (US minus CS).

#### Progressive ratio test

This test measured the motivation of marmosets to work for reward (Figure 1C). Having been trained on the touchscreen (Supplemental methods), marmosets pressed a circular stimulus on a touchscreen monitor to receive reward (5 seconds of milkshake (∼10% w/v banana Nesquik in whole milk) with a proceeding 0.5sec beep sound), with the number of required responses increasing across trials. The point at which marmosets stopped responding was termed the breakpoint. The required number of responses incremented by 1 for the first 9 trials, with this increment doubling every nine trials (Figure 1D). Sessions terminated either after 30minutes or after two minutes of inactivity.

Measures used in this test included:

- percent change in overall responses
- percent change in overall response rate

Manipulation data was always compared to the previous day’s mock infusion data; thereby taking into account any variability of these measures between weeks.

- percentage of time spent licking the spout during non-rewarded periods
- post-reinforcement pause (PRP); the latency to respond to the stimulus immediately after a reward.
- average response rate per trial (RRPT); the number of responses made over the time between the first and last response.

To calculate the average PRP and RRPT for each subject the trial number of the shortest session of each subject across all manipulations was selected as cut-off point.

#### Sucrose preference test

Sucrose preference test was used to measure the impact of scACC-25 manipulations on marmosets’ consummatory and preference behaviour, similar to Alexander and colleagues[6], using 6% sucrose solution (w/v, Sigma-Aldrich). They were presented with identical plastic bottles, one containing water and the other containing sucrose solution. Once marmosets achieved a stable sucrose preference of ∼80-90% over two sessions, the experimental manipulations took place.

Measurements included

- total sucrose consumption (g),
- total water consumption (g),
- ratio of sucrose consumption over total consumption (sucrose preference).

A detailed description of tests and a summary of their order for all subjects are described and shown in Supplemental Methods and Figure S2.

### Statistical analyses

Statistical analyses were performed using IBM SPSS Statistics (version 28). In experiment 1, a one-way repeated ANOVA (3 levels: vehicle, cortisol 1.5ng/μl, cortisol 5ng/μl), and one-sample t-test (difference score between drug and vehicle) were used. For multiple comparisons Bonferroni corrections were applied and if no correction was used it was reported. In experiment 2, two-way mixed measures ANOVAs were used in progressive ratio test for three measures of total responses, response rate and licking outside reward delivery, using drug as a within subject factor (3 levels) and water restriction as a between subject factor (2 levels). A one-way repeated measures ANOVA (3 levels: vehicle, MB, DHK) was used in progressive ratio test in measures of average response rate per trial and post-reinforcement pause, as well as in sucrose preference test for measures of sucrose preference, sucrose, and water consumption. Wherever data transformation was necessary to meet the assumptions of ANOVA, the type of transformation is reported. For post hoc analyses Sidak correction was applied for pairwise comparisons. Whenever assumption of sphericity using Mauchly’s test was not met Greenhouse-Geisser correction was applied.

### Post-mortem assessment of cannulae placement

Marmosets were premedicated with ketamine hydrochloride (0.1ml of a 100mg/mL solution, i.m.) and subsequently euthanised with sodium pentobarbital (Dolethal; 1mL of a 200mg/ml solution, i.v.). Following perfusion with 0.1M phosphate-buffered saline (PBS; Sigma-Aldrich) and 10% formalin solution, the brain was sectioned and stained with cresyl violet for cannulae placement confirmation. Histological assessment confirmed that all subjects had cannula placements in scACC-25 (Figure 2 and detailed in Supplemental Methods).

**Figure 2.**
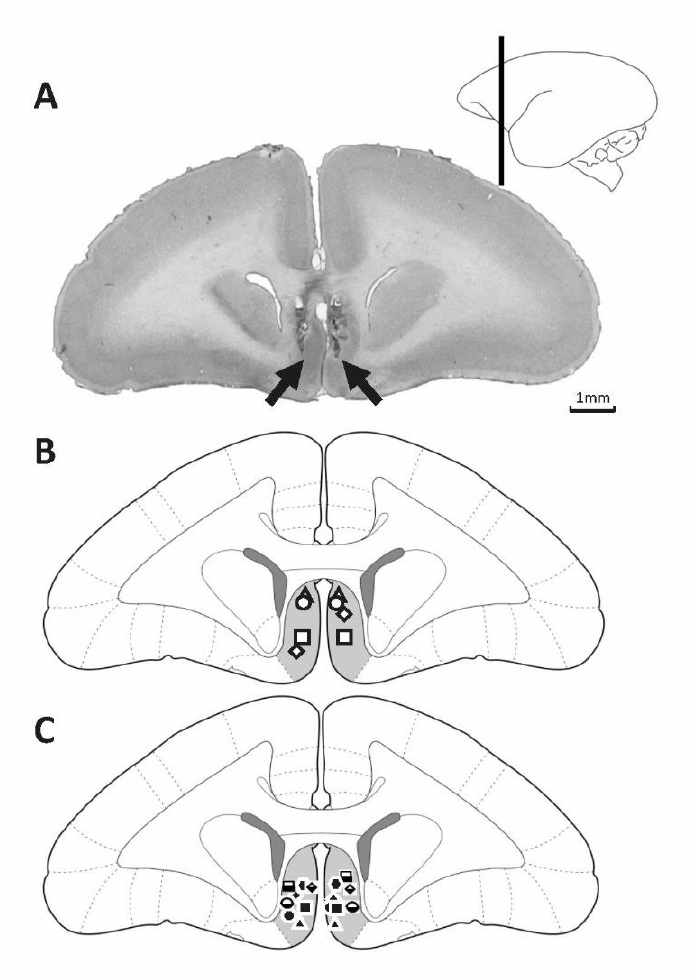
Histological verification of scACC-25 cannulation. A. Example of a cresyl stained coronal section with infusion site indicated by black arrows. B and C depict schematics indicating the confirmed infusion placements of subjects in experiments 1 and 2 respectively, within scACC-25 (light grey shading) at AP +13.80[21]. The range of infusion placements lay between AP 13.00-13.80.

## Results

### Experiment 1a: Central infusion of cortisol increased anxiety-like behaviour in a Human Intruder test

Infusion of cortisol into scACC-25 dose dependently heightened reactivity to a human intruder as revealed by an increase in the overall EFA-derived anxiety score (*F*_*(2,6)*_*=7*.*64, p=0*.*023, η*^*2*^*=0*.*717*). The increase in anxiety-like behaviour was only observed with cortisol 5ng/*μ*l (Figure 3A; *t*_*(3)*_*=3*.*26, p=0*.*047 (not corrected), CI=[0*.*01, 0*.*79]*). This was a consequence primarily of an increase in avoidance of the intruder shown by their moving further back (Figure 3B; *t*_*(3)*_*=3*.*28, p=0*.*046 (not corrected), CI=[0*.*71, 46*.*81]*) and higher (Figure 3C; *t*_*(3)*_*=4*.*89, p=0*.*016, CI=[3*.*22, 15*.*23]*) in the cage. Cortisol 1.5ng/μl did not impact any measure in this test (*EFA score p=0*.*963; TSB%, p=0*.*620; height, p=0*.*964*).

**Figure 3.**
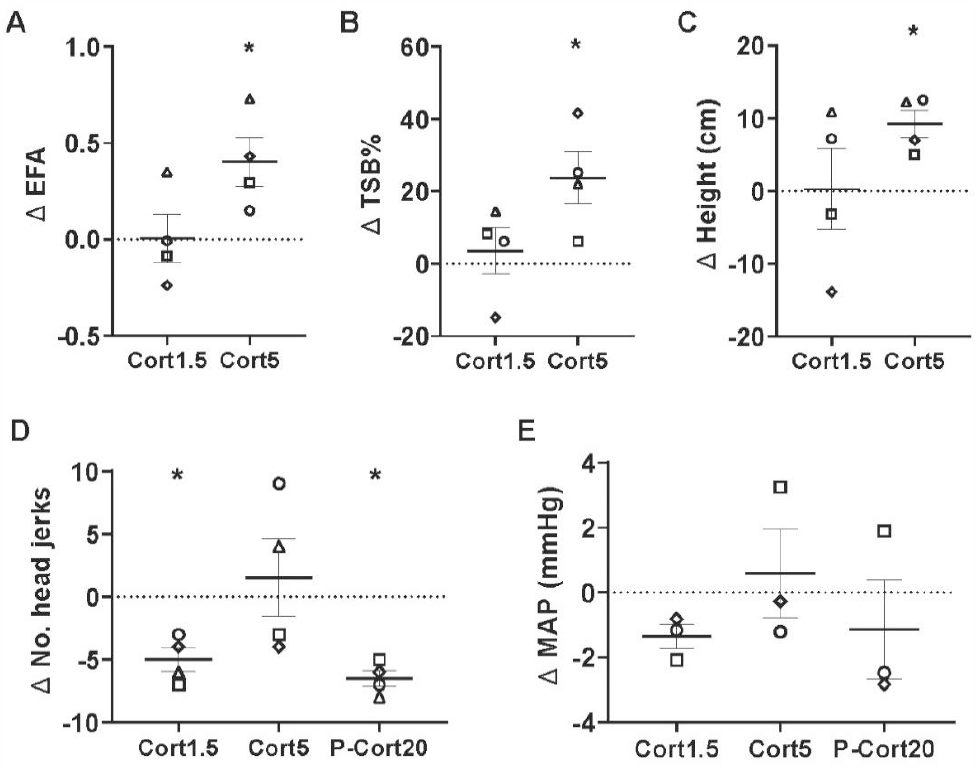
The effects of scACC-25 cortisol infusion on responsivity to uncertain threat and anticipatory arousal. A. Responsivity to an uncertain threat in the form of a human intruder was increased by scACC-25 infusion of Cort5 with an increase in the EFA score, whilst Cort1.5 had no effect. Of the factors contributing to this EFA increase were B. the time spent at the back of the cage (TSB%) and C. average height. Data are displayed as mean ± SEM error, *p<0.05. D. Infusion of cortisol into scACC-25 decreased CS+ directed head jerk behaviour at 1.5ng/*μ*l (Cort1.5) but not 5ng/*μ*l (Cort5) relative to saline. Peripheral injection of 20mg/kg cortisol also decreased this behaviour (P-Cort20). E. Neither central or peripheral administration of cortisol impacted anticipatory cardiovascular arousal, with CS-directed MAP being unaffected.

### Experiment 1b: Cortisol infusion into scACC-25 reduced anticipatory behavioural arousal in an appetitive Pavlovian test

Infusion of 1.5ng/*μ*l cortisol (*t*_*(3)*_*=-5*.*48, p=0*.*024, CI=[-7*.*91, -2*.*09]*), but not 5ng/*μ*l (*t*_*(3)*_*= 0*.*49, p=0*.*658, CI = [-8*.*27, 11*.*27]*) into scACC-25 reduced anticipatory behavioural arousal to the CS+ (Figure 3D,E), with neither dose impacting cardiovascular arousal (*1*.*5ng/μl: p=0*.*069, 5ng/μl: p=0*.*707*). Dampened behavioural arousal to the CS+ was also achieved by a peripheral injection of 20mg/kg cortisol (*t*_*(3)*_*=-10*.*07, p=0*.*002, CI=[-8*.*55, -0*.*45]*) with cardiovascular arousal similarly unaffected (*p=0*.*532*). Importantly, both CS- and US+ mediated responses were unaffected by either dose (*CS-1*.*5ng/μl: p=0*.*718, 5ng/μl: p=0*.*718; US+ 1*.*5ng/μl: p=0*.*467; 5ng/μl: p=0*.*324*) consistent with previous findings with overactivation of scACC-25[6].

### Experiment 2a: Opposing effects of scACC-25 inactivation and overactivation on motivation

Overall responses on the PR test were affected by drug manipulation (*F*_*(2,10)*_*=45*.*78, p<0*.*001, η*^*2*^*=0*.*88, cube-root transformed*), where MB-induced inactivation of scACC-25 increased (*p=0*.*048*) whilst DHK-induced overactivation of scACC-25 reduced (*p=0*.*011*) responding (Figure 4A), with the between manipulation comparison also significant (*p<0*.*001*).

**Figure 4.**
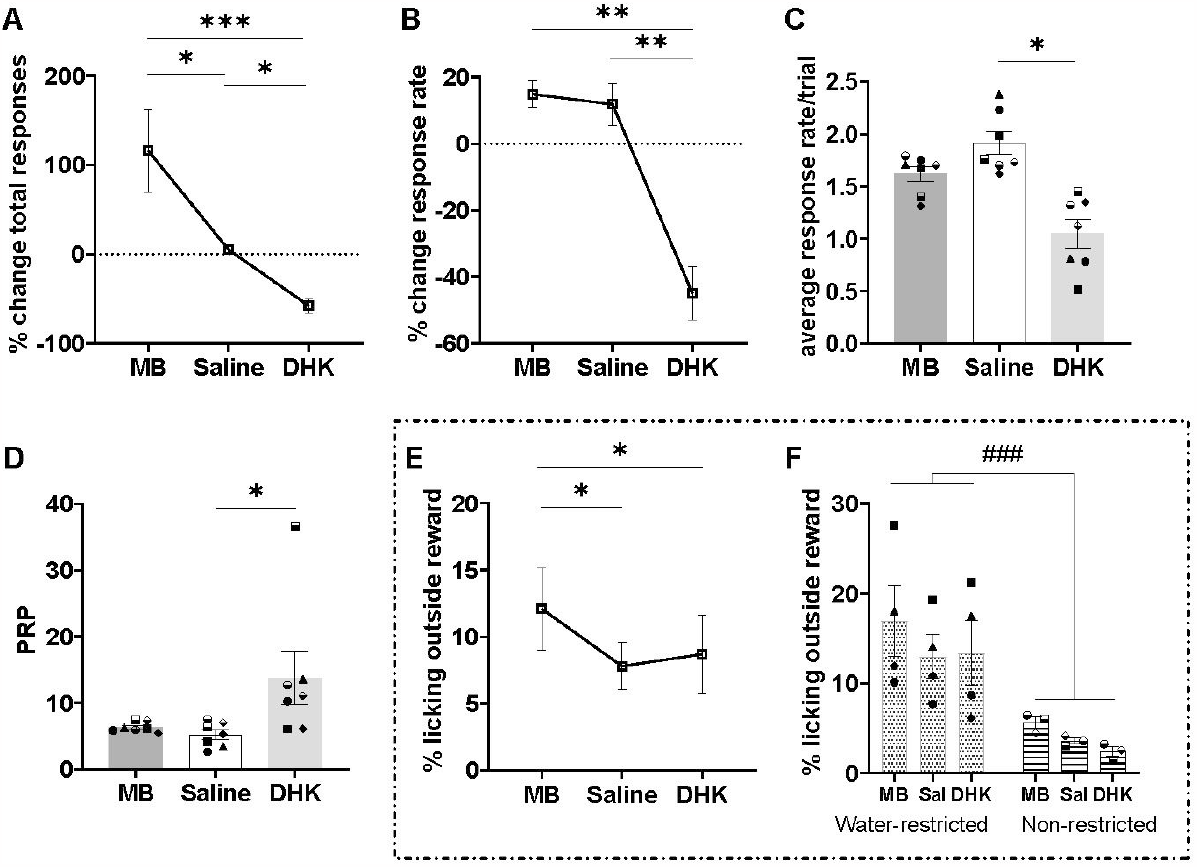
The effect of scACC-25 interventions on appetitive motivation in the progressive ratio test. A. Inactivation (MB) and overactivation (DHK) of scACC-25 led to increases and decreases in overall responses, respectively. B. DHK infusion into scACC-25 reduced overall response rate compared to saline and MB. C. Average response rate per trial was decreased by DHK compared to saline. D. DHK increased the post reinforcement pause (PRP) compared to saline. E. Inactivation of scACC-25 led to an increase in the amount of time spent licking the spout outside of reward delivery when compared to saline and DHK. F. Water restriction increased the time spent licking outside reward delivery regardless of the effect of drug manipulation. Data are displayed as mean ± SEM error, ^*^p<0.05, ^**^p<0.01 and ^***^p<0.001.

These effects were accompanied by changes in response rate (*F*_*(2,12)*_*=38*.*54, p<0*.*001, η*^*2*^*=0*.*92*) with a reduction following overactivation compared to saline (*p=0*.*002*; Figure 4B), whilst inactivation had no effect compared to saline (*p=0*.*994*). Analysis of the two components that shape response rate, average response rate per trial (*RRPT; F*_*(2,12)*_*=12*.*36, p=0*.*001, η*^*2*^*=0*.*67*), and post-reinforcement pause (*PRP; F*_*(2,12)*_*=10*.*15, p=0*.*003, η*^*2*^*=0*.*63, log-transformed*) revealed that overactivation slowed both RRPT (*p=0*.*026*) and PRP (*p=0*.*032*), whilst MB-induced inactivation was without effect compared to saline (*RRPT, p=0*.*067; PRP, p=0*.*30*, Figure 4C,D).

Inactivation saw an increase in licking outside of the reward delivery period (*F*_*(2,10)*_*=12*.*99, p=0*.*002, η*^*2*^*=0*.*72, log-transformed*; Figure 4E,F). Inactivation increased ‘off target’ licking compared to both saline (*p=0*.*023*), and overactivation (*p=0*.*025*), whereas, overactivation itself had no significant effect (*p=0*.*540*). Water restriction in general led to increased licking outside reward delivery with a significant main effect of water schedule (*F*_*(1,5)*_*=20*.*51, p=0*.*006, η*^*2*^ *=0*.*80, log-transformed*) but it did not impact overall responses or response rate (*F<1* for all analyses).

### Experiment 2b: Inactivation of scACC-25 increased, whereas overactivation decreased, sucrose consumption

There was no drug effect on sucrose preference (*F*_*(1*.*06,4*.*23)*_*= 3*.*44, p=0*.*133, η*^*2*^*=0*.*46;* Figure 5A), however, inactivation and overactivation of scACC-25 had opposing effects on sucrose consumption (*F*_*(2,8)*_*=21*.*72, p<0*.*001, η*^*2*^*=0*.*84;* Figure 5B). Inactivation increased (*F*_*(1,4)*_*=13*.*90, p=0*.*20, η*^*2*^*=0*.*78 – Simple Contrast*), and overactivation decreased (*F*_*(1,4)*_*=10*.*79, p=0*.*030, η*^*2*^*=0*.*73*) sucrose consumption, whilst neither manipulation impacted water intake (*F<1*; Figure 5C).

**Figure 5.**
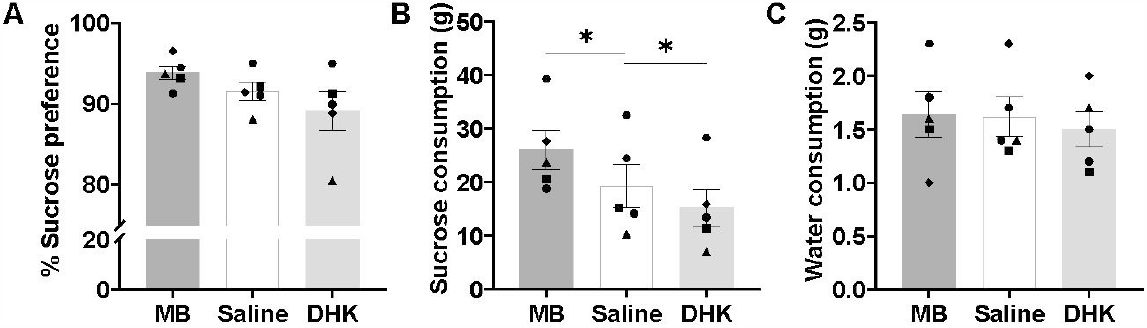
The effect of scACC-25 interventions on consummatory behaviour. A. Inactivation and overactivation of scACC-25 had no effect on sucrose preference. B. Inactivation of scACC-25 increased consumption of sucrose solution compared to saline whereas DHK decreased it. C. Neither drug manipulation had any effect on water consumption. Data are displayed as mean ± SEM error, ^*^p<0.05.

## Discussion

The present study addressed two questions. The first was whether at least some of the actions of the stress hormone cortisol were mediated via the activation of scACC-25. Results of experiment 1 showed that direct elevation of the stress hormone, cortisol, within scACC-25, resulted in a pattern of behavioural changes that mirror those seen following scACC-25 overactivation[6]. There was both a dampening of behavioural anticipatory appetitive arousal, as well as an increase in responsivity to uncertain threat. A comparable dampening of behavioural arousal was also seen following systemic administration of cortisol. The second question was regarding the extent to which scACC-25 regulated appetitive behaviours. Unlike the previously reported lack of effect of inactivation of scACC-25 on Pavlovian anticipatory arousal[6], here inactivation did impact motivated responding (experiment 2). Specifically, inactivation increased the number of responses they made for reward and increased their consummatory behaviour. The opposing effects seen following scACC-25 overactivation, namely decreased responding for reward and decreased sucrose consumption highlights the bidirectional effects of these manipulations.

### Cortisol infused locally into scACC-25 mirrors the behavioural effects of scACC-25 overactivation

Since heightened reactivity to uncertain threat and blunted Pavlovian anticipatory appetitive arousal is seen following infusions of either the glutamate transporter blocker DHK or cortisol into scACC-25 it supports the hypothesis that some of the effects of stress-induced cortisol release are dependent on scACC-25 activation. There appeared to be differential sensitivity of scACC-25 to the two doses of cortisol whereby blunting of anticipatory appetitive behavioural responses was induced by the low dose, 1.5ng/μl, whilst only the higher dose of 5ng/μl heightened responsivity to uncertain threat. When compared to systemic administration of 20mg/kg cortisol comparable effects on reward responsivity were seen, although in both cases, peripheral or central cortisol failed to significantly reduce cardiovascular arousal during this period. Whilst overall similarity in the pattern of effects is consistent with the hypothesis that cortisol acts to heighten scACC-25 activity it is unclear why behavioural anticipatory arousal is more sensitive to cortisol compared to cardiovascular anticipatory responses. This may relate to pharmacological differences between the studies, with DHK heightening activity broadly across scACC-25, whilst cortisol may impact mineralocorticoid or glucocorticoid signalling on specific scACC-25 neurons that mediate behaviour related processes rather than those involved in the central regulation of cardiovascular activity.

The timing of these effects were relatively rapid, being observed within an 8-minute window likely reflecting the fast acting, nongenomic effects of cortisol. The literature, overall, suggests that nongenomic actions of cortisol emerge within 5 minutes in brain regions such as the hippocampus[22]. Rapid non-genomic effects of cortisol were also likely to be responsible for the systemic cortisol effects in which the pre-treatment time was 15 minutes. Certainly, intramuscular injections of cortisol in humans have been shown to alter fMRI activity as early as 7 minutes, disappearing by 20 to 27 minutes[23]. In contrast, genomic actions typically start to emerge after one hour when the nongenomic effects have disappeared[24]. Recent evidence suggests that these rapid non-genomic effects of glucocorticoids, such as cortisol, may be through membrane-bound mineralocorticoid receptors inducing shifts in pyramidal neuron excitability[25]. Therefore, with the high concentration of mineralocorticoid receptors located in scACC-25 of non-human primates[13], increases in circulating cortisol, such as during periods of acute stress, may rapidly modulate behaviour via this region.

### Bidirectional effects of scACC-25 interventions on response breakpoint

The bidirectional effects of scACC-25 manipulation on motivation appear distinct from previously reported unidirectional effects of overactivation on Pavlovian appetitive arousal. Ceiling effects are unlikely to explain the lack of effect of inactivation on the latter since inactivation of neighbouring area 14 enhances behavioural and cardiovascular appetitive arousal in the same test[17]. It is worth noting that the bidirectional effects seen on breaking point of the PR test were not seen on other measures of performance. Overactivation reduced overall response rate, not only increasing the latency between individual responses leading up to a reward, but also increasing the PRP. However, inactivation had no such effects on latencies but did instead increase non-rewarded licking behaviour, suggestive of an increase in reward directed Pavlovian responses. Whether these contrasting effects reflect distinct underlying mechanisms is unclear. Due to very low levels of licking outside of reward at baseline, this measure could not have been reduced following overactivation and so bi-directional effects could not have been seen. In contrast, instrumental responses were not so high as to prevent bi-directional effects from being seen.

A likely mediator of these instrumental and Pavlovian effects is the nucleus accumbens (NAc) which receives a major input from scACC-25[26]. The NAc has been implicated in Pavlovian approach responses[27] and previous findings from our laboratory have identified the scACC-25 to NAc core pathway as a mediator of the blunting of Pavlovian appetitive behavioural and cardiovascular arousal induced by scACC-25 overactivation[28]. The NAc has also been implicated in motivated instrumental responding, with response rate, in particular, associated with perceived effort values[29] and may relate to dopamine levels[30,31]. Of particular relevance to scACC-25 is the finding in rodents that infralimbic cortex (ILc) modulates NAc dopamine firing activity, with inactivation heightening and overactivation reducing VTA activity[32]. Whilst the functional homology of scACC-25 and ILc is far from clear, with opposing effects having been reported for the regulation of threat[15], there is greater correspondence between scACC-25 and ILc with respect to the regulation of rewarded behaviour[32,33]. Thus, the reductions in response rate observed following scACC-25 overactivation in the current study may be the result of increased effort mediated through the accumbens dopamine pathway.

Consumption on the other hand is related to opioid signalling within the ventral striatum[34]. Specifically, μ-opioid receptors in the NAc shell are associated with licking and feeding behaviour[35–38] and microinjection of a *μ*-opioid agonist into the NAc shell increases “liking” of sucrose solution in rodents i.e. enhancing hedonic capacity[39,40]. In the present study scACC-25 inactivation increased food-oriented responses in the progressive ratio test but also increased sucrose consumption in the sucrose preference test. Since scACC-25 projects to both NAc subregions[41] then the effects of inactivation on consummatory responses may be mediated through alternative pathways to the NAc that mediate opioid signalling. Whilst scACC-25 overactivation did not impact licking behaviour on progressive ratio it did reduce sucrose consumption. This contrasts with our previous results where overactivation had no such effect, albeit with a higher concentration of sucrose (10%) than used here (6%)[6]. The sucrose concentration markedly alters the threshold for animals to engage with the reward[36] therefore the most likely explanation for this discrepancy is the higher concentration produced a ceiling effect, with the lower sucrose concentration here unmasking the consummatory effects of scACC-25 overactivation. Thus, an increase in the overall numbers of responses following scACC-25 inactivation on the PR test may be the result of increased reward value as distinct from an increase in response effort that likely mediates scACC-25 overactivation effects on response rate.

## Conclusion

These findings indicate scACC-25 is an important regulator of responsivity to threat and reward. They provide important insight into the functions of this region and its relevance to our understanding of the stress-related disorders of depression and anxiety. Increased activation of scACC-25 likely accompanies the stress induced by the presence of threat and, through actions of cortisol on glucocorticoid and mineralocorticoid receptors in scACC-25, enhances behaviours that increase avoidance of the threat. Since, scACC-25 overactivation also blunts consummatory, anticipatory and motivated behaviours for reward, as does scACC-25 infusions of cortisol, it is likely that these act as adaptive responses reducing the competition between approach and avoidance responses in times of threat. However, scACC-25 appears also to be important for providing tonic inhibitory control of motivated appetitive behaviours even in the absence of explicit threat, as shown by the marked increase in responding for reward, both consummatory and instrumental, when scACC-25 was inactivated. Some of these effects may be mediated through scACC-25 projections onto distinct neurochemical circuits within the NAc, which may be revealed in future studies. What is clear is that the balance of motivated behaviours to avoid threat and approach reward is in part dependent upon activity in scACC-25 and that alterations in tonic levels of activity within scACC-25, as may occur during stress, can alter that balance.

## Acknowledgements

We would like to thank Mrs Gemma Cockcroft for their assistance with histology. We would also like to thank Professor Stafford Lightman for discussions related to intra-cortical levels of cortisol. We would also like to thank Dr Kevin Mulvihill and Dr Shaun Quah for their experimental support throughout the early stages of the Covid-19 pandemic.

## Author contributions

Conceptualization – ACR, CMW, RBT; Methodology – ACR, CMW, RBT, LM; Experimentation – CMW, RBT, LM; Manuscript preparation – ACR, CMW, RBT; Manuscript editing – ACR, CMW, RBT.

