## Supplemental Methods for "Subcallosal area 25: its responsivity to the stress hormone cortisol and its opposing effects on appetitive motivation in marmosets"

#### Subjects

The room temperature and humidity were tightly controlled, with a 12-h light-dark schedule (7am lights on). For a summary of subjects used in the experiments described in the manuscript, see Table S1. All marmosets were naïve at the start of Experiment 1. Four marmosets were naïve (s1-s4) in Experiment 2. The remaining four had all previously received infusions to either activate or inactivate scACC-25 (s5[1]; s6-8[2]). Data on scACC-25 overactivation on appetitive motivation has been reported previously[2]; although only total responses were presented.

Marmosets were housed in cages (280 x 120 x 98cm) containing a nest box and a variety of environmental enrichments such as suspended ladders, wooden branches and ropes (described extensively in Alexander and colleagues[3]). Marmosets were provided water ad libitum except for s1-4 during the progressive ratio testing phase where water bottles were removed at 17:30 Sunday-Thursday, and returned at 12:00 daily. Marmosets were provided with a modified diet on Sunday-Thursday consisting of MP.E1 primate diet (Special Diet Services, UK) and two carrot slices, otherwise, their diet was supplemented with fruit, rusk, malt loaf, and eggs.

| Subject and symbol | | sex | History | Experiment 1 | | Experiment 2 | |
|---|---|---|---|---|---|---|---|
| | | | | Appetitive Pavlovian Discrimination | Human Intruder Test | Progressive Ratio Test | Sucrose Preference Test |
| *1* | ● | *M* | | | | ✓ | ✓ |
| *2* | ▲ | *F* | | | | ✓ | ✓ |
| *3* | ◆ | *M* | | | | ✓ | ✓ |
| *4* | ■ | *F* | | | | ✓ | ✓ |
| *5* | ⬣ | *M* | + | | | | ✓ |
| *6* | ◒ | *F* | + | | | ✓ * | |
| *7* | ⬙ | *F* | + | | | ✓ * | |
| *8* | ⬓ | *M* | + | | | ✓ * | |
| *1* | △ | *M* | | ✓ | ✓ | | |
| *2* | ◇ | *F* | | ✓ | ✓ | | |
| *3* | ○ | *M* | | ✓ | ✓ | | |
| *4* | □ | *F* | | ✓ | ✓ | | |

**Table S1.** Summary of subjects, their symbol, prior experimental history, and experiments in which they participated. * DHK effect on overall responses for these animals reported previously in Alexander and colleagues[2].

**Surgical procedures:**

After inducing anaesthesia marmosets were intubated and anaesthesia was maintained by 2.0–2.5% isoflurane gas in 0.3 L/min of oxygen. Using a pulse oximeter capnograph (Microcap Handheld Capnograph) respiration, heart rate (HR), oxygen saturation, and carbon dioxide blood levels were monitored throughout all surgeries. Body temperature was also monitored using a rectal temperature probe (TES-1319 K-type digital thermometer) and maintained throughout surgery via a heat mat. Postoperative analgesia was administered using meloxicam (0.1 mL of 1.5 mg/mL suspension orally, Metacam, Boehringer Ingelheim) for three consecutive days following surgery.

*Implantation of intracerebral cannulae into area 25*

Following sedation and intubation subjects were placed in a stereotaxic frame designed for marmosets (David Kopf). Two depth checks were carried out at two coordinates of +17.5 anteroposterior (AP), −1.5 lateromedial (LM), and +14.0 [AP], -1.0 [LM]. If not within the respective depth ranges of 5.8-6.8mm and 8.9-9.3mm[4,5], then the AP coordinates were adjusted in situ. Once within the range, through small holes drilled in the skull, cannulae were implanted into area 25 (C235G-1.4, 7 mm below pedestal; +14.0 AP; ±0.7 LM). The cannulae were fixed in place with skull screws, Super Bond adhesive, and dental acrylic (Paladur). Dexamethasone (0.18 mL of 3.3 mg/mL, IM, Wockhardt UK ltd) was administered immediately prior to the end of surgery to limit swelling. A weekly cleaning of the implant with sterile dummy cannulae and caps replaced.

*Implantation of the telemetry probe*

Antimicrobial liquid medicine enrofloxacin (Baytril; 0.2ml of a 2.5% solution) was administered orally 24 hours prior to surgery. Following sedation and intubation as described above, an incision was made down the midline of the animals' abdomen. Pressure was applied on the descending aorta to occlude blood for no more than three minutes. The end of the telemetry probe (HD-S10; Data Sciences International) was then inserted into the aorta at the downstream of the pressure point, and the probe was sutured in place within the abdomen.

**Drug treatment**

Sterile drug infusions into scACC-25 were carried out while an experienced helper held the marmosets gently in their hands. The procedure was also performed under sterile conditions where all the surfaces including the guide and the cement mount around the guide were cleaned with 70% isopropyl alcohol (Alcotip, Universal). Drugs were delivered bilaterally via injectors (Plastic One, C235I/SPC) connected to 10µl Hamilton syringes using PTFE tubing. A pump delivered a constant rate of 0.5µl/min for DHK (6.25nmol/µl; Tocris, UK) and two doses of cortisol (1.5 and 5ng/µl diluted in 0.9% saline; Hydrocortisone hemisuccinate, Sigma-Aldrich), and a rate of 0.25µl/min for the muscimol/baclofen cocktail (0.1mM muscimol/1.0mM baclofen; Sigma-Aldrich) for a total duration of 2 minutes. All infusions were allowed to diffuse for one minute whilst the injectors remained in place. After replacing dummy cannulae and caps, marmosets were returned to their home cage for the pretreatment period of 8 minutes for cortisol, 10 minutes for DHK, and 25 minutes for mus/bac before being carried to and placed in the testing apparatus via a transparent Perspex carry-box by the experimenter. For vehicle conditions the same procedures were implemented with saline solution instead of the drugs.

Subcutaneous injection of 20mg/kg cortisol (or saline as control) was administered with a pre-treatment time of 15 minutes.

| **Drug Vehicle** | **Mechanism** | **Route** | **Dose** | **Pre-treatment time** |
|---|---|---|---|---|
| *Cortisol Hemisuccinate Vehicle: saline Experiment 1* | *GR agonist* | *Central infusion Area 25* | *1.5, 5 ng/µl Rate of 0.5µl/min for 2 minutes* | *8 minutes* |
| *Cortisol Hemisuccinate Vehicle: saline Experiment 1* | *GR agonist* | *Systemic s.c. injection* | *20 mg/Kg 0.8 ml/Kg* | *15 minutes* |
| *Dihydrokainic acid (DHK) Vehicle: saline Experiment 2* | *$EAAT_2$ antagonist* | *Central infusion Area 25* | *1.35 µg/µl Rate of 0.5 µl/min for 2 minutes* | *10 minutes* |
| *Muscimol/baclofen (MB) Vehicle: saline Experiment 2* | *$GABA_A$ /$GABA_B$ receptor agonist* | *Central infusion Area 25* | *11.4 ng/µl muscimol 0.214 µg/µl baclofen Rate of 0.25 µl/min for 2 minutes* | *25 minutes* |

**Table S2.** Details of drugs used in these experiments, their route of administration, dose and pre-treatment time, which is the time interval between completion of infusion and the time test commences. After infusion is completed the injectors remain in place for an extra minute to allow for adequate diffusion. An $EAAT_2$ (Excitatory amino acid transporter-2) inhibitor increases overall levels of glutamate at the synapse by reducing the amount of glutamate taken up by its transporter.

**Behavioural testing apparatus**

Both the progressive ratio and appetitive Pavlovian tests were carried out in a sound-attenuated apparatus, whilst the human intruder and sucrose preference tests were carried out in the top right quadrant of the home cage, where the subject was divided away from the partner. The testing apparatus was fitted with a house light (3 W), speakers and three cameras connected to Power Director software (Cyberlink). A telemetry receiver (Physiotel, Data Sciences International) was located under the floor of the chamber, with data recorded by Spike 2 software (Cambridge Electronic Design).

*Human intruder test*

The human intruder test was used to measure the intolerance of marmosets to uncertainty under experimental manipulations (described in detail in Quah and colleagues[6]. The test consists of three phases: separation, intruder, and recovery. Marmosets are initially restricted to the top right quadrant for 8 minutes (separation phase). An experimenter wearing a realistic latex human mask (Greyland Film, UK) and standard lab attire enters the room and stands in front of the home cage for two minutes and maintains constant eye contact with the subject (Intruder phase). The intruder then leaves and the marmosets behaviour is recorded for another 5 minutes (recovery phase). The subject's behaviour on the test, was recorded by a GoPro Hero 5 video camera on a tripod. Behaviour was scored offline (JWatcher software), measuring the time subjects spends in each zone (Figure S1). From this positioning, the percentage time moving, head and body bobs and vocalisations (recorded by a shotgun microphone) were compiled into an exploratory factor analysis (EFA) score[6].

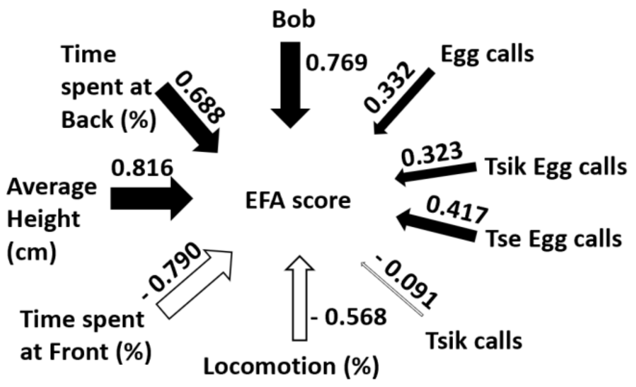


**Figure S1.** The factor loading of each measure used in the EFA score which is derived from an exploratory factor analysis from a cohort of 171 subjects.

*Appetitive Pavlovian test*

For a full description of this test and the testing apparatus refer to Alexander and colleagues[3] (Figure 1). Briefly, prior to conditioning, all marmosets were habituated to the apparatus and reaching for freely available high incentive food (several pieces of marshmallow) in the food box. They were then habituated to the sight and sound of the food box door opening revealing reward. Once the subjects stopped showing a mild startle response (i.e., rearing and jumping) to the opening of the door, and started consuming marshmallow within 40 s of its opening, they were advanced to conditioning sessions where two distinct tones were played to cue reward (box full of marshmallow) and no reward, (empty food box) respectlvely. A biweekly testing schedule was implemented Monday to Friday with a total of 5 rewarded trials. Minor variations of this schedule were used to prevent anticipation of rewarded trials. Each session started with a 60 second acclimatisation period and a variable inter-trial interval (ITI, 70-110s) in two trial sessions in which experimental manipulations were performed (Figure 1B).

*Progressive ratio test*

Marmosets were first trained to touch the screen for reward, as described in full in Alexander and colleagues[2]. Briefly, subjects were first familiarised with the delivery of banana milkshake from a spout in the testing apparatus, and then trained to respond to a stimulus presented on a touchscreen for reward. The stimulus first was a green rectangle across the width of the screen and following successful touching to gain reward, it was changed to a square stimulus presented centrally and then to the left or right of the licker[7]. Once marmosets were reliably and accurately responding to this stimulus for a reward the stimulus was changed to a white circle presented at a fixed location (the animal's preferred side). Fixed ratio (FR) response schedules were then used to train the subjects to make repeated responses for reward with schedule progressing from FR1 to FR7 after which they were moved to progressive ratio schedule[8].

Measures used in the progressive ratio test included percent change in overall responses or overall response rate, with manipulation data compared to the previous day where they received a mock infusion. This is to take into account any variability of these measures between weeks. Other measures included the percentage of time spent licking the spout during non-rewarded periods and the post-reinforcement pause (PRP), which is the latency to initiate a response from when the stimulus is presented at the start of each trial[9]. To calculate the average PRP for each subject the trial number of the shortest session of each subject across all manipulations was selected as cut-off point. Average response rate per trial was also calculated based on the number of responses made in each trial over the time between first and last response. This measure was then averaged over the shortest trial number of each subject across all manipulations.

*Sucrose preference test*

Sucrose preference test was used to measure the impact of scACC-25 manipulations on marmosets consummatory and preference behaviour, similar to Alexander and colleagues[2], using 6% sucrose solution (w/v, Sigma-Aldrich) via transparent plastic bottles. Marmosets were first habituated to the sucrose solution by introducing two identical 6% sucrose solution bottles in the upper right quadrant of the cage for 48h. Subsequently, a marmoset was separated from their cage mate by dividing them off into the top right quadrant of the home cage with the nest box and usual water bottle removed. They were then presented with the same two bottles, one bottle containing water and the other containing sucrose solution.

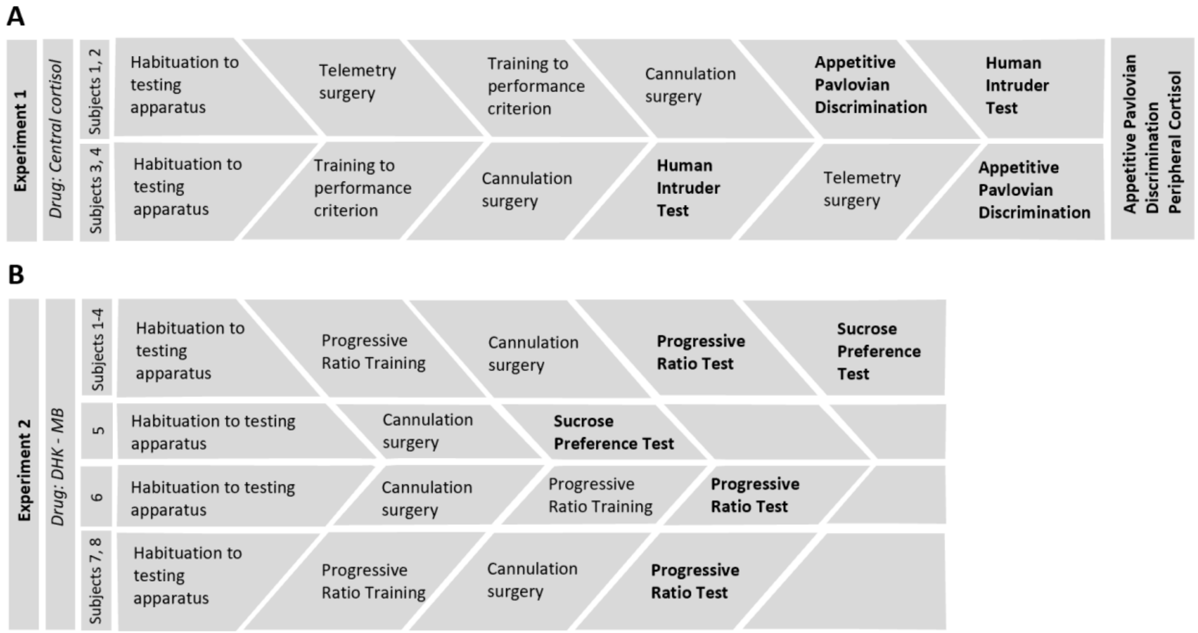


**Figure S2.** Flow chart demonstrating the order of experiments carried out in A. Experiment 1 and B. Experiment 2. In Experiment 1 (n=4) , half the marmosets (subjects 1 and 2) were tested on appetitive Pavlovian discrimination before the human intruder and vice versa for the other half (subjects 3 and 4). All subjects then proceeded to appetitive Pavlovian test and peripheral cortisol manipulation. In experiment 2, seven out of eight marmosets were tested on progressive ratio, four of which were also tested on sucrose preference. One animal was only tested on sucrose preference.

### Post-mortem assessment of cannulae placement

Marmosets were premedicated with ketamine hydrochloride (0.1ml of a 100mg/mL solution, i.m.) and subsequently euthanised with sodium pentobarbital (Dolethal; 1 mL of a 200mg/ml solution, i.v.). Following this, transcardial perfusion of 500ml 0.1M phosphate-buffered saline

(PBS; Sigma-Aldrich) was conducted followed by 500ml of 10% formalin solution. The brain was then extracted and placed in 10% formalin solution for 24 hours, then 0.01M PBS-azide solution for 48 hours and finally 30% sucrose for 72 hours. The brains were then cut into 40µm coronal sections and mounted on gelatin-coated slides and stained with cresyl violet to confirm cannula placement. Sections were then visualised and photographed using a M205FA stereo microscope (Leica, UK). Histological assessment confirmed that subjects successfully had cannula placement into area 25 (Figure 2).
